# Differential spatiotemporal representations along the hippocampal long axis in humans

**DOI:** 10.1101/179655

**Authors:** Iva K. Brunec, Buddhika Bellana, Jason D. Ozubko, Vincent Man, Jessica Robin, Zhong-Xu Liu, Cheryl Grady, R. Shayna Rosenbaum, Gordon Winocur, Morgan D. Barense, Morris Moscovitch

**Affiliations:** Department of Psychology, University of Toronto; Rotman Research Institute, Baycrest; Department of Psychology, SUNY Geneseo; Department of Psychology, York University; Department of Psychology, Trent University; Department of Psychiatry, University of Toronto

## Abstract

Increased place field size and signal autocorrelation along the dorsoventral hippocampal axis in rodents are considered a fundamental aspect of hippocampal organization, yet such evidence is lacking in humans. Using fMRI, we report corresponding evidence of increasing neural similarity from posterior to anterior hippocampus (dorsoventral homologues) in humans. These findings help account for observed shifting in representational granularity, from global context (anterior) to local details (posterior), along the hippocampal axis.

---

The ability to represent the world accurately relies on simultaneous coarse and fine-grained neural coding of information, such that both gist and detail of an experience are preserved. The longitudinal axis of the hippocampus has been proposed to provide a gradient of representational granularity in spatial and episodic memory in rodents and humans [1-3, 6-8]. Rodent place cells in the ventral hippocampus exhibit significantly larger place fields and greater autocorrelation than those in the dorsal hippocampus [1, 4, 5], which may underlie a coarser and slower-changing representation of space in ventral hippocampus, and correspondingly, a more rapidly shifting, finer delineation of space in the dorsal hippocampus [1,4] (Fig. 1A).

**Figure 1.**
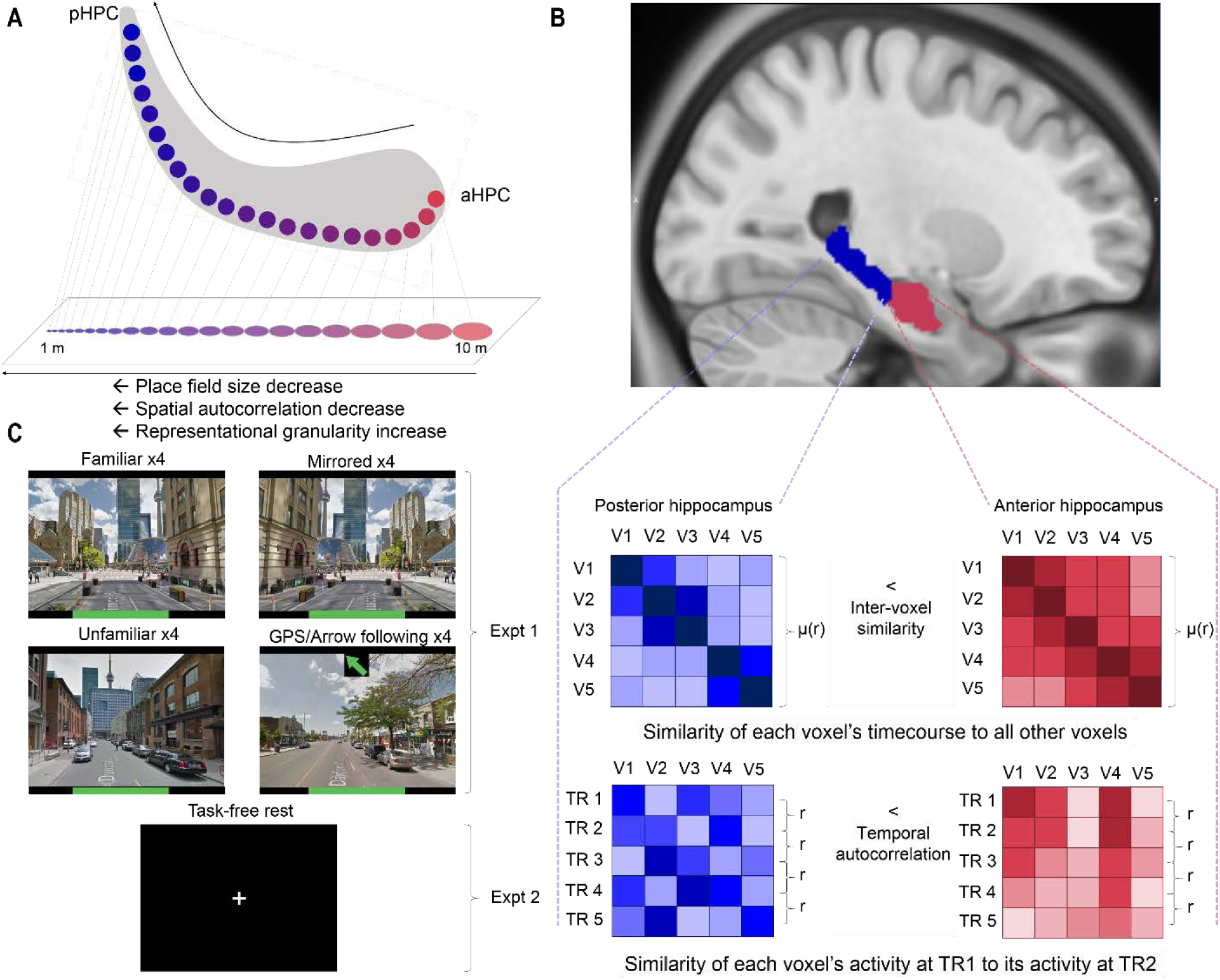
A) The hypothesized increase in granularity along the anteroposterior axis is derived from rodent research suggesting a decrease from place field size in the ventral (anterior in humans) to dorsal (posterior in humans) hippocampus. B) Anterior (red) and posterior (blue) hippocampal masks overlaid on an MNI template brain and conceptual illustration of inter-voxel similarity and temporal autocorrelation measures. Inter-voxel similarity was calculated as the mean of all unique correlations in each region, hemisphere, and condition, per participant. This is akin to measures of functional connectivity, calculated between voxels within an ROI. Temporal autocorrelation was derived by calculating pairwise correlations for all voxels in an ROI at successive timepoints (TRs). The mean of the resulting r-coefficients was calculated for each region, hemisphere, and condition, per participant. Note that the cells in the inter-voxel similarity matrix reflect voxel-wise correlations, while the cells in the temporal autocorrelation matrix reflect voxel-wise activity. C) In Experiment 1, participants navigated in 4 conditions: familiar paths, unfamiliar paths, familiar paths where the environment was left-right reversed, and GPS/arrow following on unfamiliar paths. In Experiment 2, participants rested with their eyes open and later reported how much time they spent engaging in episodic simulation (thinking about future episodes or remembering the past).

Although direct electrophysiological measurements along the entire axis are impractical in humans, evidence for place-specific cells has been reported [9-11]. It remains unclear, however, whether the dynamic cellular properties along the hippocampal long axis are comparable across humans and rodents. Recent evidence suggests that properties of cellular dynamics robustly demonstrated in rodents can be captured with fMRI in humans during spatial navigation [12] and conceptual learning [13]. It is possible that the mechanisms supporting granularity along the long axis may also be extrapolated to the scale of fMRI signal. Here, using fMRI, we provide novel evidence for the existence of separable signal properties along the human hippocampal anteroposterior axis, and relate them to navigation and episodic memory.

If fMRI voxel-wise activation patterns are influenced by the increase in granularity stemming from larger place fields in anterior hippocampus (aHPC, the homologue of ventral hippocampus in rodents) to smaller place fields in posterior hippocampus (pHPC, dorsal in rodents), their fMRI timecourses should be more similar to one another in the aHPC, where the representations would be coarser (Fig. 1A-B), than in the pHPC. In addition, based on the reported increase in autocorrelation found in rodent place cell firing along the dorsoventral axis, voxel activation patterns may also show more autocorrelation over successive time points in aHPC than in pHPC (Fig. 1B).

To test the hypothesis that dynamics of hippocampal activity in human fMRI reflect differences in granularity along the hippocampal long axis, we calculated 1) correlations between voxel timecourses within aHPC and pHPC (i.e., similarity in activity across voxels within an ROI; *inter-voxel similarity*) and 2) autocorrelation of the fMRI signal for individual voxels within aHPC and pHPC (i.e., similarity in each voxel’s activity across successive time points; *temporal autocorrelation*). We predicted that both types of neural similarity would be higher in aHPC, compared to pHPC, corresponding to a coarser level of representation and more slowly changing signal over time. We examined these measures during virtual reality navigation under different conditions to investigate, for the first time, neural dynamics underlying representational granularity along the hippocampal axis in humans (Fig. 1C).

Participants in the navigation task (N = 19; see Supplementary Methods) traversed routes in a familiar virtualized environment (Toronto, Canada), using images from Google Street View (Fig. 1C). Each participant traversed 16 routes while undergoing fMRI. Automated Anatomical Labeling atlas (AAL) hippocampal masks from each hemisphere were divided into anterior and posterior portions based on the location of the uncal apex (MNI space y = -21) [2]. We then calculated 1) inter-voxel similarity and 2) temporal autocorrelation (Fig. 1B).

To test for differences in inter-voxel similarity along the hippocampal axis, we conducted a 2 (anterior vs posterior axis) x 2 (hemisphere) x 4 (condition) repeated-measures ANOVA. Inter-voxel similarity was significantly greater in aHPC, relative to pHPC (F(1, 18) = 50.06, p < .001; Fig. 2A), with a significant main effect of hemisphere (F(1, 18) = 39.02, p < .001), and a significant interaction between axis and hemisphere (F(1, 18) = 16.12, p < .001), driven by a larger difference between aHPC and pHPC in the left, relative to the right hippocampus (Fig. S1). There was no significant main effect of condition (F(3, 54) = 1.31, p = .280). No other interactions were significant (all p-values ≥ .31). To visualize differences in inter-voxel similarity along the hippocampal axis with finer spatial resolution, bilateral hippocampal masks were split into 6 sub-segments with inter-voxel similarity calculated for each segment separately (Fig. 2D; for detailed analyses see Supplementary Materials). Inter-voxel similarity was again consistently higher in segments anterior to the uncal apex relative to posterior segments (significant main effect of axis; F(5, 90) = 19.98, p < .001), supporting the initial dichotomy. Interestingly, the data did not appear to show a smooth linear gradient for this measure.

**Figure 2:**
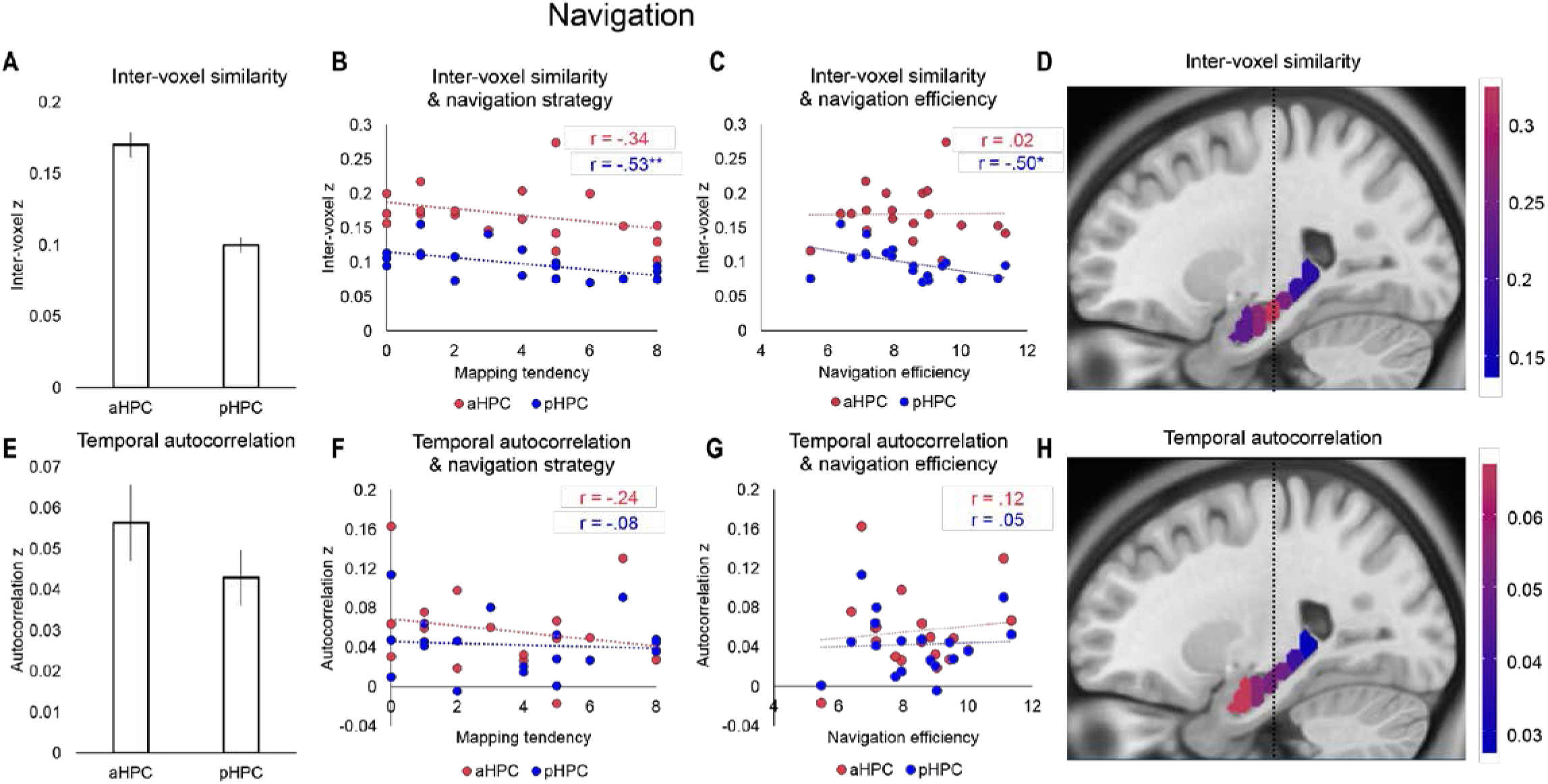
A) Inter-voxel similarity was significantly greater in aHPC, relative to pHPC, during navigation (r-coefficients were z-transformed before comparisons were made). For individual matrices for all participants, see Fig. S4. B) Inter-voxel similarity in pHPC significantly negatively correlated with the tendency to use map-based navigation strategies (relative to landmark-based strategies). C) Inter-voxel similarity in pHPC, but not aHPC, significantly negatively correlated with in-scan navigational performance/efficiency. D) Mean inter-voxel similarity along the left hippocampal axis, split into 6 segments (MNI space, x = -25). The dotted line indicates the location of the uncal apex (MNI space y = -21), which was used to guide the anterior/posterior division. E) Autocorrelation was significantly greater in aHPC, relative to pHPC during navigation. F) No significant relationships between temporal autocorrelation and navigation tendency were found. G) Unlike inter-voxel similarity, autocorrelation showed no significant relationship with in-scan navigational efficiency. H) Mean temporal autocorrelation in each of the 6 long axis segments. For further analyses, please refer to Figs. S6-S8. Error bars are s.e.m.

To explore differences in temporal autocorrelation, an identical 2 (axis) x 2 (hemisphere) x 4 (condition) repeated-measures ANOVA was conducted on this measure. Temporal autocorrelation was also significantly greater in aHPC, relative to pHPC (F(1, 18) = 7.79, p = .012; Fig. 2E). There was a significant main effect of condition, such that temporal autocorrelation scaled up with navigational demand (F(3, 54) = 21.54, p < .001; see Fig. S2). There was no significant main effect of hemisphere (F(1, 18) = .288, p = .598), but a significant interaction between axis and hemisphere (F(1, 18) = 8.44, p = .009; see Fig. S2), as well as a significant interaction between axis, condition, and hemisphere (F(3, 54) = 3.04, p = .037; see Fig. S2). No other interactions were significant (all p-values > .05; see Supplementary Materials). Furthermore, temporal autocorrelation appeared to have a smooth gradient with the most autocorrelated signal at the head of the aHPC and least in the most posterior segments of the pHPC (Fig. 2H; main effect of axis F(5, 90) = 16.73, p < .001; for detailed analyses, see Supplementary Materials).

Participants’ self-reported propensity to rely on map-based strategies in navigation was significantly negatively correlated with inter-voxel similarity in pHPC (r = -.535, p = .018), but not aHPC (r = -.341, p = .153; Fig. 2B, full questionnaire in Appendix A). Lower similarity among hippocampal voxels may imply finer-grained coding, as the signal is less redundant and can thus contain more complex information. Reduced similarity may support a greater propensity to use more complex, map-based navigational strategies that enable fine-grained spatial representations and discrimination [17]. To support this notion, we also found a significant negative correlation between pHPC inter-voxel similarity and navigational efficiency in the navigation task (r = -.503, p = .028), but no significant relationship in aHPC (r = .015, p = .951; Fig. 2C). Navigational efficiency was calculated as the average decrease in Euclidean distance to the goal between steps (greater difference therefore indicates greater navigational efficiency). Correspondingly, there was a significant relationship between self-reported mapping propensity and navigational efficiency (r = .560, p = .013), indicating that participants with greater mapping propensity navigated more efficiently towards the goal.

There was no significant relationship between map-based navigation and temporal autocorrelation in aHPC (r = -.240, p = .322) or pHPC (r = -.081, p = .742; Fig. 2F), and no significant relationship between navigational efficiency and temporal autocorrelation in aHPC (r = .116, p = .636) or pHPC (r = .051, p = .836; Fig. 2G).

Inter-voxel similarity and temporal autocorrelation were not significantly correlated in aHPC (r = .102, p = .677) or pHPC (r = .280, p = .245), suggesting that they capture separate aspects of hippocampal coding.

To establish whether these findings were task-driven (i.e., navigation-dependent) or whether they reflect a task-independent property of hippocampal dynamics, we examined the same measures in a resting state dataset carried out on a separate group of 20 participants [14]. Resting state has no explicit demand on behaviour, although continuous dynamics can still be measured, making it an ideal contrast to navigation. The aHPC and pHPC masks used on participants in the resting state analyses were identical to those used in the navigation study. We conducted 2 (axis) x 2 (hemisphere) repeated-measures ANOVAs on both inter-voxel similarity and temporal autocorrelation.

As in navigation, resting inter-voxel similarity was significantly greater in aHPC, relative to pHPC (F(1, 19) = 36.37, p < .001; Fig. 3A). There was no significant main effect of hemisphere (F(1, 19) = .317, p = .580), but a significant axis by hemisphere interaction (F(1, 19) = 4.93, p = .039), again driven by a larger difference between the left aHPC and pHPC, relative to the right (Fig. S1).

**Figure 3:**
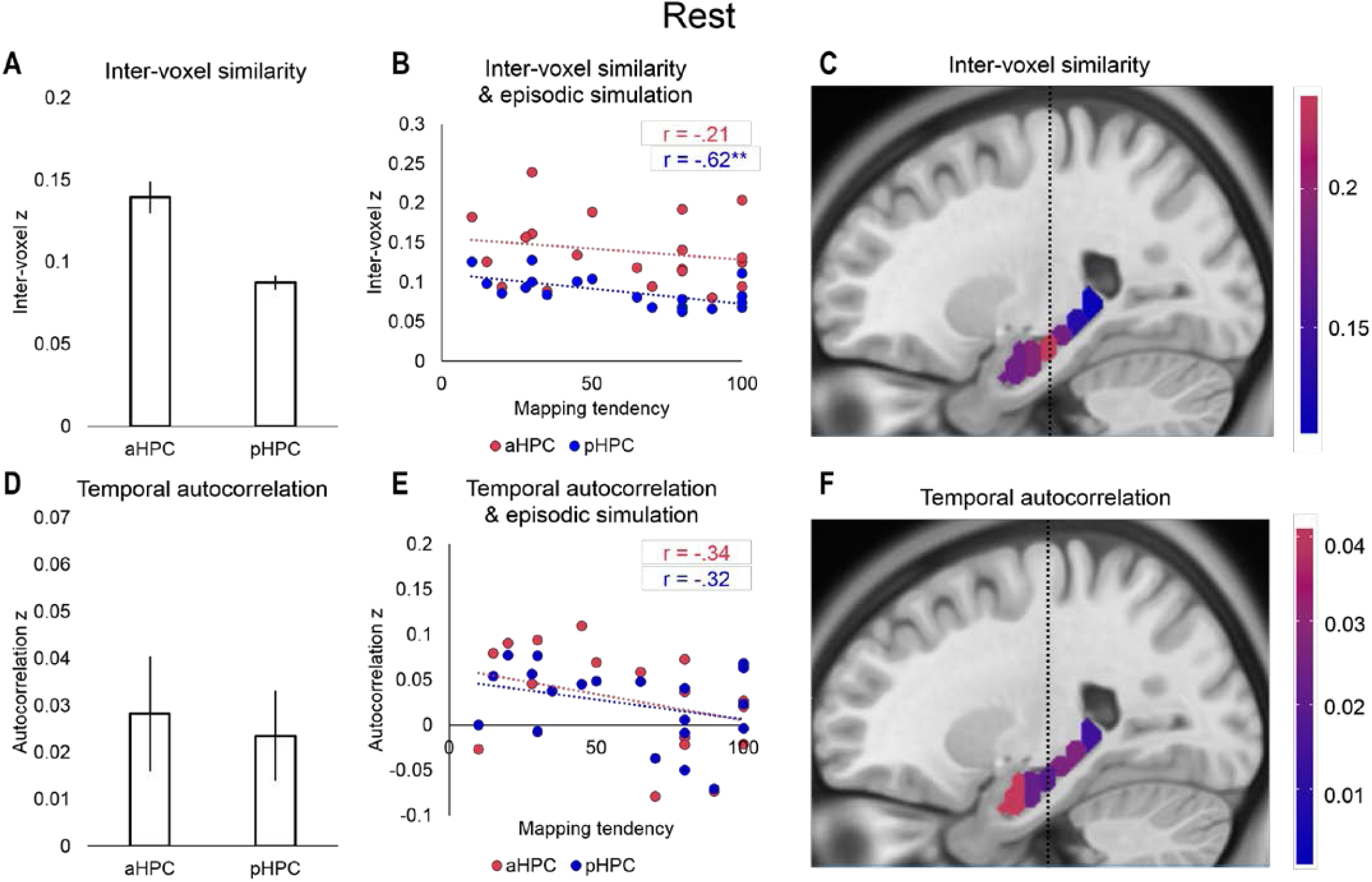
A) Inter-voxel similarity was significantly greater in aHPC, relative to pHPC during rest. For individual matrices for all participants, see Fig. S5. B) Inter-voxel similarity in pHPC significantly negatively correlated with self-reported time spent in episodic simulation (thinking about the past or simulating the future). C) Mean inter-voxel similarity along the left hippocampal axis, split into 6 segments. The dotted line indicates the location of the uncal apex (MNI space y = -21), which was used to guide the anterior/posterior division. D) Autocorrelation was not significantly different between aHPC and pHPC at rest. E) No significant relationships between temporal autocorrelation and episodic simulation were found. F) Mean temporal autocorrelation in each of the 6 long axis segments. For further analyses, please refer to Figs. S6-S8. Error bars are s.e.m.

In contrast to navigation, there was no significant difference in resting temporal autocorrelation between aHPC and pHPC (F(1, 19) = .654, p = .429; Fig. 3D), nor was there a significant main effect of hemisphere (F(1, 19) = 2.88, p = .106), or axis by hemisphere interaction (F(1, 19) = .074, p = .788).

These data suggest that some aspects of the different signal properties in aHPC and pHPC are fundamental and present in task and resting state, but appear to be modulated by the processing of incoming information or mental simulation.

While resting state scans involve no explicit processing demands, the degree to which participants engage in typically hippocampusl-dependent thought, such as thinking about the past or the future, may vary [14]. Participants’ self-reported in-scan episodic simulation (i.e., the percentage of the resting scan where they were either remembering past, or imagining future episodes) was significantly negatively correlated with inter-voxel similarity in pHPC (r = -.618, p = .004), but not aHPC (r = -.205, p = .385), akin to the relation found for map-based strategies in the navigation data (Fig. 3B). There was no significant relationship between temporal autocorrelation and self-reported episodic simulation in either aHPC (r = -.336, p = .148) or pHPC (r = -.318, p = .172; Fig. 3E).

Inter-voxel similarity and temporal autocorrelation were again not significantly correlated in aHPC (r = .386, p = .093) or pHPC (r = .268, p = .253) at rest. Inter-voxel similarity and temporal autocorrelation also differ in their similarity across aHPC and pHPC (Fig. S3).

Even though the timepoint-to-timepoint temporal autocorrelation in rest was not significant, calculating the same measure on finer spatial and temporal scales revealed the presence of a gradient (Fig. S7 & S8).

Based on the above evidence suggesting anterior-posterior differences in granularity, we sought to determine whether our findings reflect a general anteroposterior organizing principle not restricted to the hippocampus, but rather merely corresponds to a characteristic of fMRI signal properties in this region of the brain. An ideal control region would be a proximal extrahippocampal region that is highly functionally connected to the hippocampus, providing a conservative test of whether the spatiotemporal gradient extends to regions with coupled dynamics. To isolate such a region, we investigated the whole-brain functional connectivity pattern during rest for aHPC and pHPC ROIs bilaterally. For each of the four hippocampal ROIs (left and right aHPC and pHPC), we calculated the mean timecourse across voxels within the ROI, and computed voxelwise correlation values between the mean timecourse of each ROI and the timecourse of each non-hippocampal voxel in the brain.

Group-level analyses revealed that all four voxels with peak hippocampal connectivity were within the parahippocampal gyrus (PHG; Fig. 4A), a region frequently involved in spatial processing [19]. The similar long-axis orientation of the hippocampus and PHG makes the latter an ideal control to probe whether signal properties along the anteroposterior axis of the brain might account for the results presented here. Therefore, we divided AAL PHG masks into anterior and posterior portions (at the midpoint along the y-axis), per hemisphere, and re-ran inter-voxel similarity and temporal autocorrelation analyses on the resting state data.

**Figure 4:**
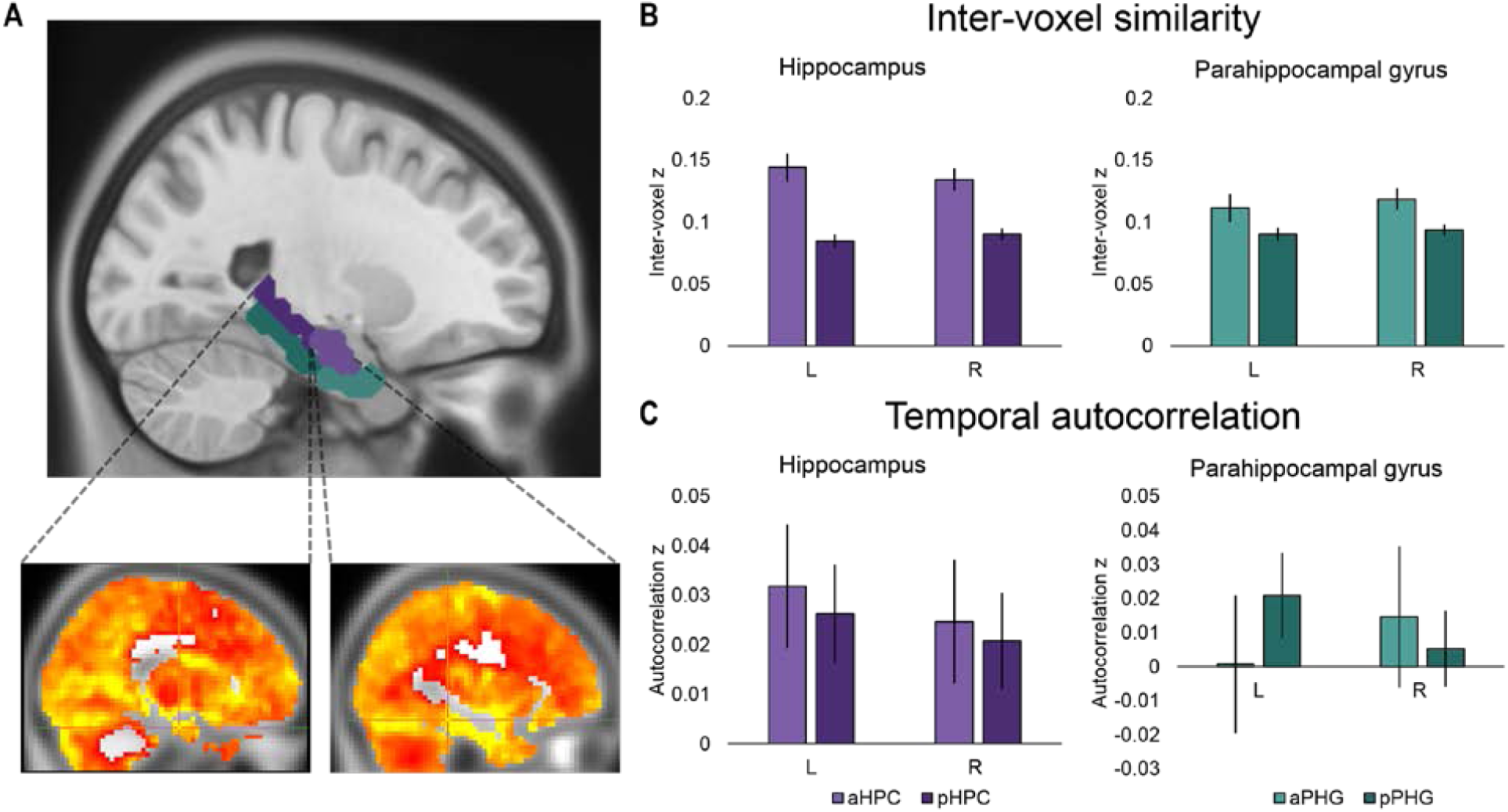
A) Left aHPC (light purple) and pHPC (dark purple) AAL masks, overlaid on an MNI template brain, with the anterior and posterior sections of the parahippocampal gyrus (PHG) below displayed in green. Displayed below are the functional connectivity maps with crosshairs indicating voxels with highest connectivity values with the left pHPC (x = 32, y = 34, z = 9) and aHPC (x = 17, y = 26, z = 12). B) The difference between anterior and posterior inter-voxel similarity was greater in the hippocampus, relative to the PHG. C) Temporal autocorrelation revealed different patterns in the left and right PHG, relative to the left and right hippocampus. Error bars are s.e.m.

If there is an extrahippocampal anteroposterior difference in granularity, it should be equally present in the hippocampus and PHG. We therefore directly compared aHPC and pHPC to anterior and posterior PHG (aPHG, pPHG) using 2 (region) x 2 (axis) x 2 (hemisphere) repeated-measures ANOVAs on both measures of inter-voxel similarity and temporal autocorrelation.

The analysis of inter-voxel similarity revealed a significant main effect of axis (F(1, 19) = 36.03, p < .001), a significant main effect of region (F(1, 19) = 12.15, p = .002), and a significant interaction between axis and region, suggesting the anteroposterior difference was greater in the hippocampus, relative to the PHG (F(1, 19) = 16.99, p < .001; Fig. 4B). There was no significant main effect of hemisphere (F(1, 19) = .250, p = .623), no significant interaction between axis and hemisphere (F(1, 19) = 1.05, p = .318), no significant interaction between region and hemisphere (F(1, 19) = 2.36, p = .141), but a significant three-way interaction between axis, region, and hemisphere (F(1, 19) = 8.75, p = .008; Fig. 4B). A two-way ANOVA comparing aPHG and pPHG bilaterally revealed a significant main effect of axis (F(1, 19) = 18.74, p < .001), with no significant axis by hemisphere interaction and no significant main effect of hemisphere (both p > .230).

Temporal autocorrelation revealed no significant main effect of axis (F(1, 19) = .002, p = .962) and no significant interaction between axis and region (F(1, 19) = 1.20, p = .288), but a main effect of region that approached significance (F(1, 19) = 4.27, p = .053). There was again no significant main effect of hemisphere (F(1, 19) = 1.16, p = .295), but a significant three-way interaction between axis, region, and hemisphere (F(1, 19) = 18.28, p < .001; Fig. 4C). The same analysis performed on navigation data similarly revealed robust differences between the hippocampus and PHG, bolstering the argument that the gradations of representational granularity presented here are fundamental to the hippocampal anteroposterior axis (Fig. S9).

Using two approaches to investigate human hippocampal activity during navigation and rest, we show robust and reliable differences in neural dynamics in the anterior and posterior hippocampus. Neural activity patterns in pHPC were not only more variable at each moment in time (inter-voxel similarity), but also shifted more rapidly between successive time points (temporal autocorrelation), relative to aHPC during navigation. During task-free rest, the difference in inter-voxel similarity between aHPC and pHPC remained significant, but no significant difference was found in temporal autocorrelation between aHPC and pHPC (though see Supplementary Materials Fig. S8). The same consistent pattern was not found in the parahippocampal gyrus, a region in close anatomical proximity to and with strong functional connectivity to the hippocampus, suggesting the pattern of findings reported here is a fundamental and distinctive organizing property of the hippocampus.

The difference in spatiotemporal representational granularity along the hippocampal axis may therefore rely on underlying neural dynamics which in aHPC, relative to pHPC, produce 1) greater similarity between individual voxel timecourses and 2) slower changes in activity. Critically, these measures were not significantly correlated, suggesting they are indicative of different aspects of the hippocampal code. These results also suggest that the cellular mechanisms observed in rodents [4, 5] may have downstream effects which scale to timescales measureable in fMRI, which support the reported findings of increased representational granularity along the axis in memory [6], narratives [3, 15], and spatial navigation [8]. Greater neural similarity in aHPC supports the notion that voxels remain in the same state for a longer time than in pHPC, which may enable the retention of gist and larger areas of space indicative of global context, but be less conducive to processing local details. Interestingly, our results did not provide consistent evidence for a smooth, coarse-to-fine, linear gradient along the hippocampal long axis. Temporal autocorrelation showed a linear decrease in similarity from aHPC to pHPC, but inter-voxel similarity only provided evidence for a dichotomy, again highlighting independence between the two measures. Further treatment of this issue is warranted to better understand the organization of the hippocampal gradient, particularly with higher resolution imaging of the hippocampus.

Differences between aHPC and pHPC were also reflected in participants’ self-reported navigation and episodic simulation tendencies. Greater propensity for map-based navigation strategies was related to lower inter-voxel similarity among hippocampal voxels. Lower similarity may suggest that information is represented in a more complex, detail-rich manner. More signal complexity could afford more flexibility, consistent with the use of cognitive maps [17]. Lower inter-voxel similarity in pHPC signal also corresponded to increased self-reported episodic simulation during rest. Remembering past and imagining future episodes reliably recruits regions of the MTL [14, 18], and greater dissimilarity among hippocampal voxels may reflect increased elaboration in episodic memory [6] and simulation [20].

At a more general level, our findings suggest that modulation in the intrinsic dynamics of hippocampal neurons may underlie or support the cognitive functions mediated by them, such as aspects of spatial navigation and episodic memory and, in combination with differences in functional connections, may help account for specialization along the long axis of the hippocampus [16]. Robust differences in granularity between aHPC and pHPC were found despite the fact that our measures did not directly manipulate granularity per se. Future studies are needed to determine whether changes in representational demands of the task will be reflected in changes in granularity as assessed by inter-voxel similarity and temporal autocorrelation, or whether changes in task demands will merely determine where along the axis activation is most pronounced.

The novel evidence presented here supports a cross-species generalization of hippocampal coding properties by revealing a potential fundamental principle by which experience in the world is coded by the hippocampus. Applying the approach to processing dynamics of hippocampal neurons reported in this study to any temporally-extended fMRI dataset, opens new doors for exploring potential differential trajectories in aging and disease, and individual differences in normal memory and cognition.

## Acknowledgments

This work was supported by Canadian Institute of Health Research grant (#49566) to R.S.R., C.G., G.W., and M.M. I.K.B. is supported by an Alzheimer Society Canada doctoral award (#17-19).

